# Intestine-on-chip enhances nutrient and drug metabolism and maturation of iPSC-derived intestinal epithelial cells relative to organoids and Transwells

**DOI:** 10.1101/2024.06.30.601390

**Authors:** Renée Moerkens, Joram Mooiweer, Eline Smits, Marijn Berg, Aarón D. Ramírez-Sánchez, Rutger Modderman, Jens Puschhof, Cayetano Pleguezuelos-Manzano, Robert J. Barrett, Cisca Wijmenga, Iris H. Jonkers, Sebo Withoff

## Abstract

The human intestinal epithelial barrier is shaped by various biological and biomechanical influences such as growth factor gradients and the flow of intestinal contents. Exposure to these cues *in vitro* impacts the cell type composition and function of adult stem cell (ASC)-derived intestinal epithelial cells, but their effect on human induced pluripotent stem cell (hiPSC)-derived cells is largely unexplored. Here, we characterize and compare the cellular composition and gene expression profiles of hiPSC-derived intestinal epithelial cells exposed to various medium compositions and cultured as organoids, in Transwell and in microfluidic intestine-on-chip systems. We demonstrate that inhibition and activation of the WNT, BMP, NOTCH and MAPK pathways regulates the presence of dividing, absorptive and secretory epithelial lineages within these systems, as has been described for ASC-based systems. Upon differentiation, intestinal epithelial organoids and monolayers in Transwell systems expressed genes involved in important intestinal functions, including digestive enzymes, nutrient transporters and members of the Cytochrome P450 family implicated in drug metabolism. However, the dynamic microenvironment of the intestine-on-chip system induced the strongest upregulation of these genes, with an expression profile that suggests a more mature developmental state. Overall, these results underscore the value of hiPSC-derived intestinal epithelial cells for modeling important functions of the human intestinal epithelial barrier and facilitates the selection of relevant culture conditions for specific applications.

## Introduction

The human intestinal epithelial barrier consists of a variety of proliferative, secretory and absorptive epithelial subtypes that facilitate the digestion and transportation of nutrients while preventing the passage of harmful environmental agents and microbes. It has now become clear that these processes are subject to inter-individual differences that depend on the host’s genetic background^1,2^. This emphasizes the importance of personalized intestinal model systems for investigating processes such as digestion, drug metabolism or drug sensitivity. Human intestinal epithelial cells can be derived from adult stem cells (ASCs) found in intestinal tissue^3^ or from human induced pluripotent stem cells (hiPSCs)^4^, which are reprogrammed somatic cells^5^. In contrast to ASC-derived tissues, which rely on invasive procedures to collect human intestinal material, hiPSCs can be created from easily accessible somatic cells from blood, skin or even urine^6^. Moreover, the pluripotent character of hiPSCs and the variety of differentiation protocols now available enable the generation of multiple cell types and tissues with an identical genetic background to study cellular interactions.

The diversity and functioning of intestinal epithelial subtypes *in vitro* are impacted by exposure to biological and biomechanical cues that emulate intestinal processes, such as growth factor gradients, tissue elasticity or the shear stress imposed by intestinal motions^7–9^. To date, most studies investigating the impact of these culture conditions on intestinal epithelial cells have been performed using ASC-derived cells. The specification of intestinal epithelial lineages is well-established for ASC-derived cells and is exerted through inhibitors or activators of the WNT, BMP, NOTCH and MAPK pathways, resembling the signaling patterns over the crypt-villus axis in the human intestine^10^. This process is not as well-controlled for hiPSC-derived intestinal tissues and often depends on spontaneous lineage induction achieved through elongated periods of culturing^4^ or maturation *in vivo* in mice^11,12^. Previous studies using ASC-derived intestinal tissues in microfluidic intestine-on-chip systems have demonstrated the value of continuous fluid flow and shear stress for the induction of villus-like folds and the expression of gene families that regulate nutrient and drug metabolism and absorption, i.e. Cytochrome P450, solute carrier (SLC) transporters and digestive enzymes^9,13^. In addition to the cellular impact of these biomechanical cues, a model system’s design can be instrumental for emulating specific aspects of the intestinal epithelial barrier. Model systems with two compartments, such as Transwell and intestine-on-chip systems, allow easy access to the apical and basolateral sides of the intestinal epithelial barrier. This facilitates the application of growth factor gradients to increase epithelial diversity and enables the study of barrier function or translocation of compounds, which is challenging in organoids due to their closed lumen and heterogeneity in size and shape.

In this study, we investigate the effect of diverse culture conditions on the subtype composition, gene expression profiles and maturation status of hiPSC-derived intestinal epithelial cells. We compared three widely used model systems: intestinal organoids, Transwell systems and a commercially available intestine-on-chip system, along with different medium compositions to enrich for specific epithelial subtypes. Hereby, we provide insight into the relevant conditions and systems for modeling specific intestinal functions using hiPSC-derived intestinal epithelial cells.

## Results

### WNT, BMP, NOTCH and MAPK regulators induce specific epithelial lineages in hiPSC-derived intestinal epithelial organoids and monolayers in a Transwell system

In ASC-derived intestinal tissues, regulators of the WNT, BMP, NOTCH and MAPK pathways efficiently enrich for specific epithelial subtypes, providing methods to control the epithelial composition *in vitro*^10,14^. Here, we investigated the contribution of these pathways to epithelial fate specification in hiPSC-derived intestinal epithelial organoids and monolayers in a Transwell system.

To generate a relatively pure population of intestinal epithelial cells, we subjected intestinal organoids from three hiPSC lines to immunomagnetic EPCAM selection and multiple subsequent passaging steps during a 4 to 5-week expansion phase, reducing the EPCAM-negative cells that co-developed during the differentiation to less than 0.1% (Figure S1A). After seeding these intestinal epithelial cells in Matrigel domes or on Transwell inserts, we exposed them to an expansion medium (EM), allowing the formation of spheroids (7 days) or a polarized epithelial barrier (14 days), respectively (Figure 1A). The subsequent differentiation phase consisted of a 5-day exposure to various medium compositions that have been described to control the induction of different epithelial lineages (Figure 1A)^10,14^. In ASC-based tissues, EM (including WNT pathway activators and BMP pathway inhibitors) induces expansion of proliferating cells, while differentiation medium (DM; excluding WNT activators and BMP inhibitors) yields a general differentiation-inducing medium. To induce differentiation toward secretory lineages, we added a NOTCH inhibitor (DAPT) to DM (DM+D). To enrich for enteroendocrine cells, we added a MAPK inhibitor (PD0325901) to DM+D (DM+D+P).

**Figure 1.**
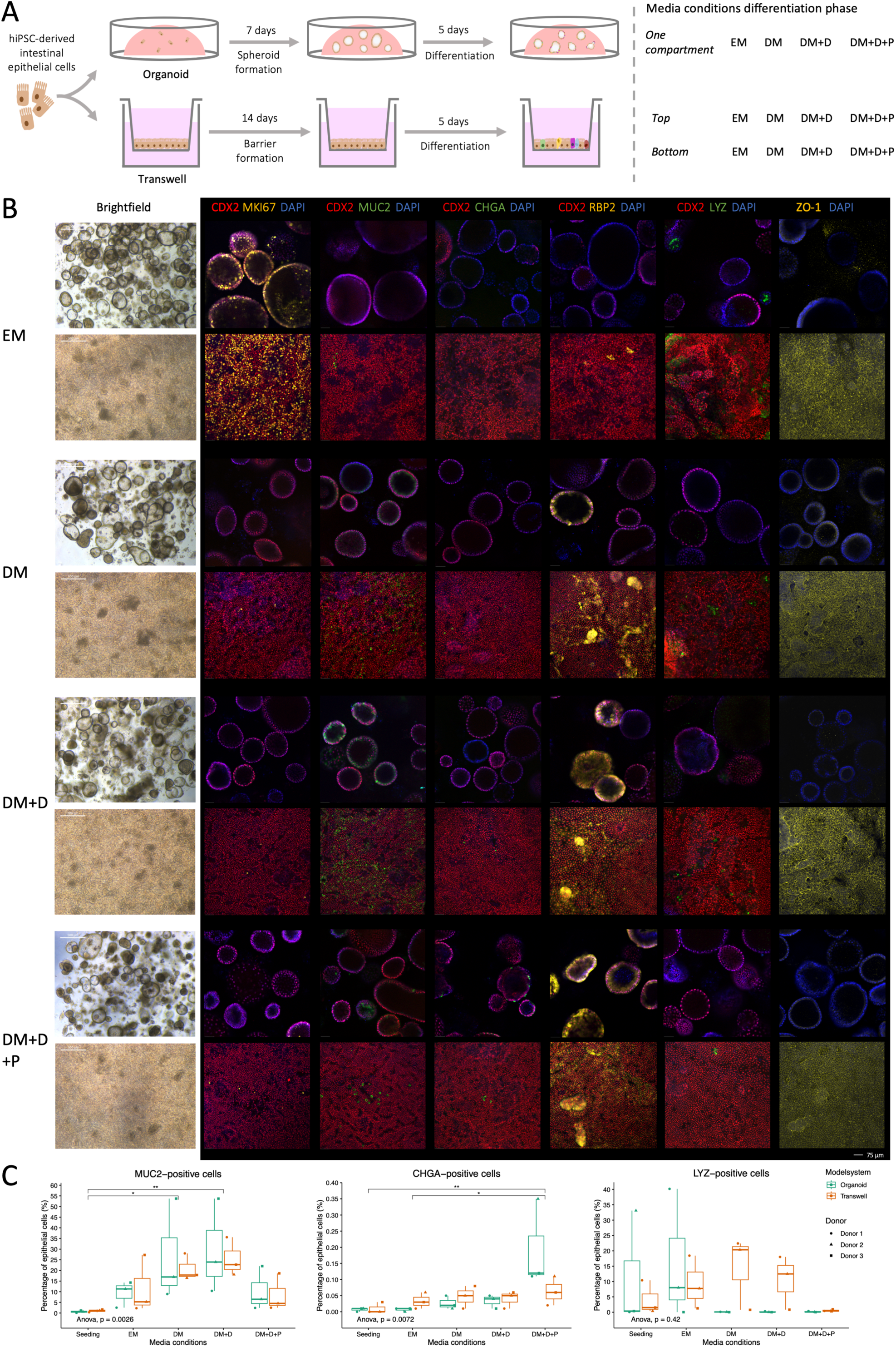
Inducing specific epithelial lineages in hiPSC-derived intestinal organoids and monolayers in a Transwell system. (A) Schematic overview of the experimental set-up. hiPSC = human induced pluripotent stem cell, EM = expansion medium, DM = differentiation medium, DM+D = DM + DAPT, DM+D+P = DM + DAPT + PD0325901. (B) Representative brightfield and immunofluorescent confocal images of hiPSC-derived intestinal epithelial organoids (upper lane) and monolayers on Transwell inserts (lower lane) stained for markers characteristic of intestinal epithelial subtypes: intestinal epithelial cells (CDX2), dividing cells (MKI67), goblet cells (MUC2), enteroendocrine cells (CHGA), enterocytes (RBP2), Paneth cells (LYZ) and tight junctions (ZO-1). (C) Quantification of cell type proportions based on flow cytometry analysis, displayed as median values of three biological replicates. P-value ≤ 0.05 (*), 0.01 (**).

Proliferative epithelial cells (MKI67-positive), likely stem cells and transit-amplifying cells, were abundant in the EM condition but scarce in the different DM conditions (Figure 1B). Transcriptional analysis comparing the EM and DM+D+P conditions also revealed that exposure to DM+D+P led to decreases in the stem cell markers *LGR5* and *SMOC2* and in the cell cycle-associated genes *MKI67*, *TOP2A*, *PCNA* and *CENPF* (Figure S1B). Immunofluorescence analysis showed that all three types of DM increased the number of RPB2-positive enterocytes, whereas very few were observed in the EM condition (Figure 1B). MUC2-positive epithelial cells, annotated as goblet cells, were present in the EM condition (organoid: 9.4% (SD 6.1), Transwell: 11.6% (SD 13.7)) but were induced to higher levels in DM (organoid: 26.5% (SD 23.9), Transwell: 20.8% (SD 6.3)) and DM+D (organoid: 29.3% (SD 22.1), Transwell: 25.5% (SD 9.1)), which is formulated to induce secretory progenitors (Figures 1B-C). The induction of MUC2-positive goblet cells was abrogated by additional MAPK inhibition in DM+D+P (organoid: 10.3% (SD 10.4), Transwell: 8.5% (SD 8.8)), in accordance with literature describing that goblet cells arise from highly-proliferating secretory progenitors (Figures 1B-C)^10^. In contrast, CHGA-positive enteroendocrine cells arise from slow-dividing secretory progenitors^10^ and were induced by the inhibition of the MAPK pathway (DM+D+P), although their numbers remained relatively low (organoid: 0.2% (SD 0.1%), Transwell: 0.06% (SD 0.05)) (Figures 1B-C). All these epithelial subtypes showed similar trends in organoids and Transwell systems, except for LYZ-positive cells (Figures 1B-C). Based on the absence of defensin gene expression, these cells were assumed to resemble the Paneth-like cell population previously identified by our group in a hiPSC-intestine-on-chip^15^. While LYZ-positive Paneth-like cells were present in all conditions except DM+D+P in Transwell systems, they were only detected in the EM condition in organoids, suggesting a differential sensitivity to the microenvironments of the organoids and Transwell systems (Figures 1B-C). The induction of cell type-specific markers, as determined on protein level (Figures 1B-C), corresponded with differences in gene expression levels of these and other markers characteristic of enterocytes, goblet cells, enteroendocrine cells and Paneth cells measured by RNA-sequencing for the EM and DM+D+P conditions (Figure S1B). The network of tight junctions between epithelial cells, as measured by the expression and localization of ZO-1, remained intact in all media conditions regardless of the induction of different epithelial subtypes (Figures 1B, S1B). Intestinal organoids were organized with the apical side projected toward the lumen, as shown by the localization of ZO-1 (Figure 1B).

Overall, in both organoids and Transwell systems, the numbers of proliferating epithelial cells were reduced and the numbers of differentiated cells were increased upon exposure to different types of DM. The removal of WNT activators and BMP inhibitors in DM was sufficient to induce goblet cells and enterocytes, while the addition of NOTCH and MAPK inhibitors (DM+D+P) was necessary to induce low levels of enteroendocrine cells. LYZ-positive Paneth-like cells were differentially induced in organoids and Transwell systems.

### Growth factor gradients in Transwell systems balance proliferating and differentiating epithelial subtypes

Next, we assessed whether a local inhibition and activation of the WNT, BMP, NOTCH and MAPK pathways could sustain both dividing and differentiated epithelial subtypes. We exposed the Transwell system to EM in the lower compartment and DM+D or DM+D+P in the upper compartment to mimic the growth factor gradients along the crypt-villus axis in the human intestine (Figure 2A).

**Figure 2.**
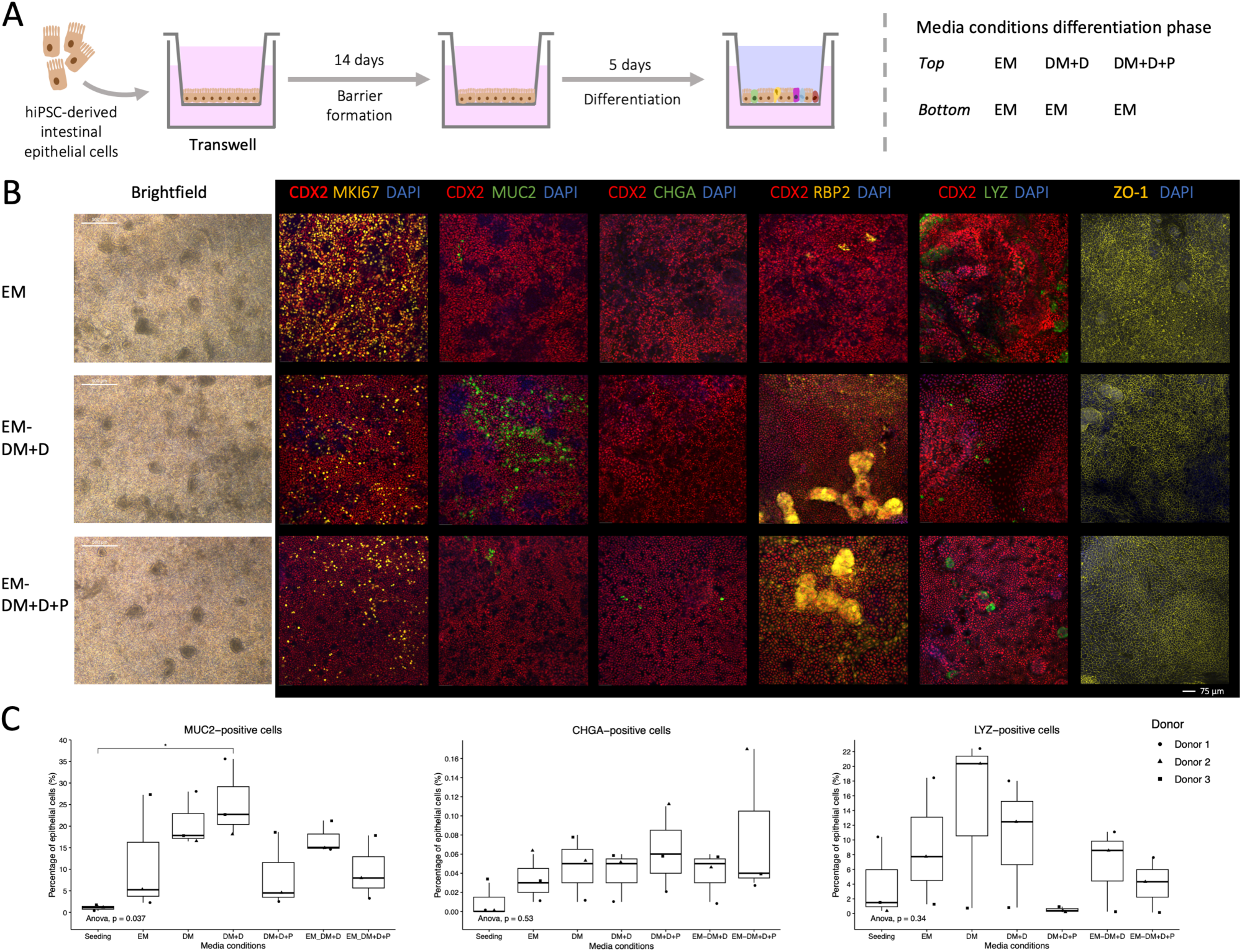
Balancing proliferating and differentiating epithelial subtypes using growth factor gradients in Transwell systems. (A) Schematic overview of the experimental set-up. hiPSC = human induced pluripotent stem cell, EM = expansion medium, DM = differentiation medium, DM+D = DM + DAPT, DM+D+P = DM + DAPT + PD0325901. (B) Representative brightfield and immunofluorescent confocal images of hiPSC-derived intestinal epithelial monolayers on Transwell inserts stained for markers characteristic of intestinal epithelial subtypes: intestinal epithelial cells (CDX2), dividing cells (MKI67), goblet cells (MUC2), enteroendocrine cells (CHGA), enterocytes (RBP2), Paneth cells (LYZ) and tight junctions (ZO-1). (C) Quantification of cell-type proportions based on flow cytometry analysis, displayed as median values of three biological replicates. P-value ≤ 0.05 (*).

In both gradient conditions, proliferating epithelial cells were sustained while differentiated epithelial subtypes were induced (Figure 2B). More RBP2-positive enterocytes were observed in both gradient conditions relative to the EM condition (Figure 2B). In concordance with observations in DM+D and DM+D+P only, MUC2-positive goblet cells were induced in EM-DM+D (17% (SD 3.72)) and CHGA-positive enteroendocrine cells were induced in EM-DM+D+P (0.08% (SD 0.08)) when compared to the EM condition (Figures 2B-C). The levels of LYZ-positive Paneth-like cells in EM-DM+D were similar to those seen in the EM condition but slightly reduced in EM-DM+D+P (Figures 2B-C). A network of tight junction protein ZO-1 was observed in both gradient conditions, suggesting the preservation of barrier integrity (Figure 2B). We further confirmed the more balanced expression of dividing and differentiated epithelial subtypes upon exposure to EM-DM+D+P relative to EM and DM+D+P on gene expression level (Figure S1B).

Strikingly, donor 3 displayed lower levels of stem cell and cell cycle genes and LYZ-positive Paneth-like cells and higher levels of markers corresponding to differentiated epithelial subtypes (e.g. MUC2) in all media conditions and in the EM condition in particular (Figures 2C, S1B). We noticed that these observations correlated with a lower expression of VIM in donor 3 compared to donors 1 and 2 (Figure S1B). VIM is a marker characteristic of mesenchymal cells that was expressed at low levels regardless of the limited number of EPCAM-negative and VIM-positive mesenchymal cells (below 0.1% upon seeding) (Figures S1A-B). It is possible that higher numbers of mesenchymal (VIM-positive) cells in donors 1 and 2 support the undifferentiated and Paneth cell state of hiPSC-derived intestinal epithelial cells and hereby prevent spontaneous differentiation.

In conclusion, gradient conditions in Transwell systems efficiently induced differentiated epithelial subtypes, including enterocytes, goblet cells and enteroendocrine cells, while preserving proliferating epithelial cells, thereby providing a more physiological and potentially sustainable condition.

### Transcriptomic analysis confirms enrichment of enterocyte- and enteroendocrine-specific functions upon exposure to DM+D+P

To obtain deeper insight into the biological processes corresponding to the changing epithelial compositions, we compared the transcriptomic profiles of hiPSC-derived intestinal epithelial cells in different medium conditions using RNA-sequencing. We used organoids grown in the EM and DM+D+P conditions for this analysis because we observed a diverse epithelial composition, including goblet cells, enteroendocrine cells and enterocytes, in the latter condition. We clustered differentially expressed genes (DEGs) between the conditions into two groups depending on whether they were upregulated in the EM or DM+D+P condition and then identified biological processes by pathway enrichment analysis (Figures 3A-C, Data S1). DEGs upregulated in the EM condition were involved in processes related to DNA replication, translation and cell division (Figures 3B-C), consistent with the high abundance of proliferating epithelial cells in the EM condition relative to DM+D+P (Figure 1B). In the DM+D+P condition, we observed the upregulation of genes associated with nutrient metabolism and transport, xenobiotic metabolism and hormone responses, reflecting the induction of enterocytes and enteroendocrine cells (Figure 3B). Specifically, we observed the induction of digestive enzymes (e.g. *ANPEP, SI, LCT, PEPD, LIPA* and *ALDOB*) and nutrient transporters (e.g. *APOB, APOA2, SLC2A2, SLC15A1, SLC5A9* and *SLC7A9*) involved in protein, carbohydrate and lipid metabolism characteristic of the human small intestine^16,17^ (Figure 3C). Many of these processes were related to lipid metabolism and synthesis of carboxylic and organic acids (Figure 3B), common by-products of small intestinal digestion^18^. We also observed increased expression of drug-metabolizing enzymes and transporters (e.g. *CYP3A4, CYP3A5, CY1A1, CYP2C9, ABCC2, UGT2A3* and *MAOB*), comprising multiple genes associated with first-pass metabolism of xenobiotic compounds in the intestine^19,20^ (Figures 3B-C). *CYP3A4*, one of the most well-studied drug-metabolizing enzymes and responsible for the conversion of more than half of all drugs^21^, showed the highest induction of all the Cytochrome P450 members identified in DM+D+P relative to EM (Figure 3C). In addition, we observed an induction of (pro-)hormones involved in gut permeability and immune activation (*CHGA*)^22,23^, gastric motility (*MLN*)^24^ and pH regulation (*SCT*)^25^ in the human intestine (Figure 3C).

**Figure 3.**
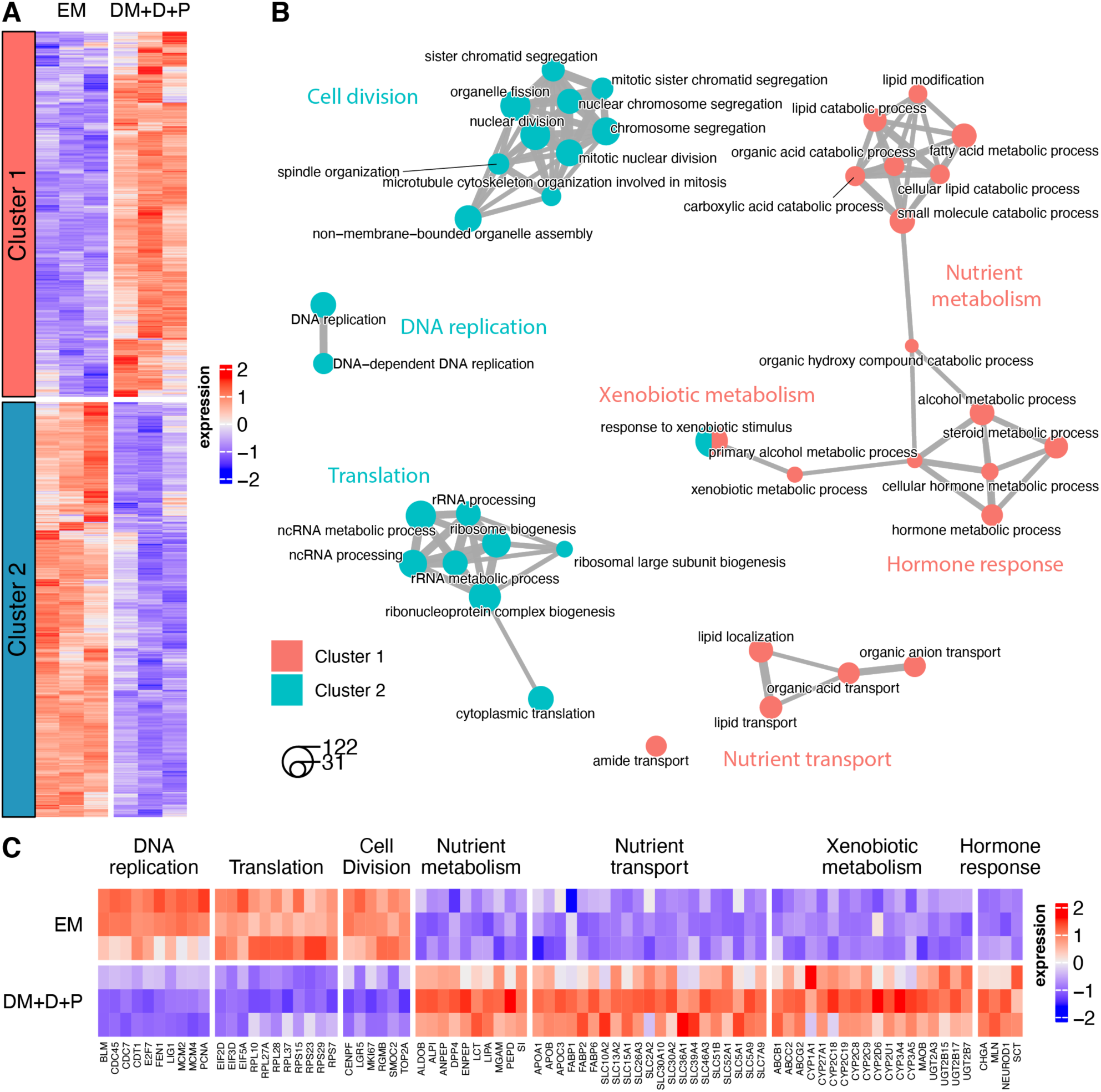
Enrichment of enterocyte- and enteroendocrine-specific functions in the DM+D+P condition in intestinal organoids. (A) Heatmap of average expression of the DEGs between medium conditions in intestinal organoids, clustered in groups of genes with similar expression profiles. Color scale represents row z-score. Columns correspond to donors, left=1, middle=2, right=3. (B) Network of enriched processes identified using the Gene Ontology: biological process database (top 20 with lowest adjusted p-value per cluster). Nodes depict enriched biological processes. Node size indicates the number of DEGs associated to the process. Edge thickness correlates with the number of DEGs shared between processes. (C) Heatmap of average expression of DEGs selected from enriched processes described in B. Color scale represents column z-score. Rows correspond to donors, upper=1, middle=2, lower=3.

The same analysis was performed using intestinal epithelial cells grown in Transwell systems and exposed to the EM, DM+D+P and gradient EM-DM+D+P conditions (Figures 4A-C, Data S1). The genes and biological processes induced by the different conditions were relatively consistent between organoids and Transwell systems and included the induction of DNA replication and cell division in the EM condition and nutrient and xenobiotic metabolism and transport as well as hormone responses in the DM+D+P condition (Figures 4B-C). The EM-DM+D+P condition in Transwell systems induced gene expression profiles intermediate between the profiles induced in the EM and DM+D+P conditions, in accordance with the presence of proliferating and mature epithelial subtypes we observed before (Figures 2B-C, 4A). Interestingly, in contrast to organoids, pathways and genes related to neurogenesis were upregulated (e.g. *TUBB2B, NCAM1, CRABP2, ETV5, NRP1* and *RGMA*) in the EM condition in Transwell systems, which might correspond to previous observations of neuron development in a hiPSC-derived intestine-on-chip exposed to the same medium^15^ (Figures 4B-C). Regardless of the limited number of EPCAM-negative cells, these mesenchymal and neural cell types might still be present at low levels and be specifically amplified in Transwell systems.

**Figure 4.**
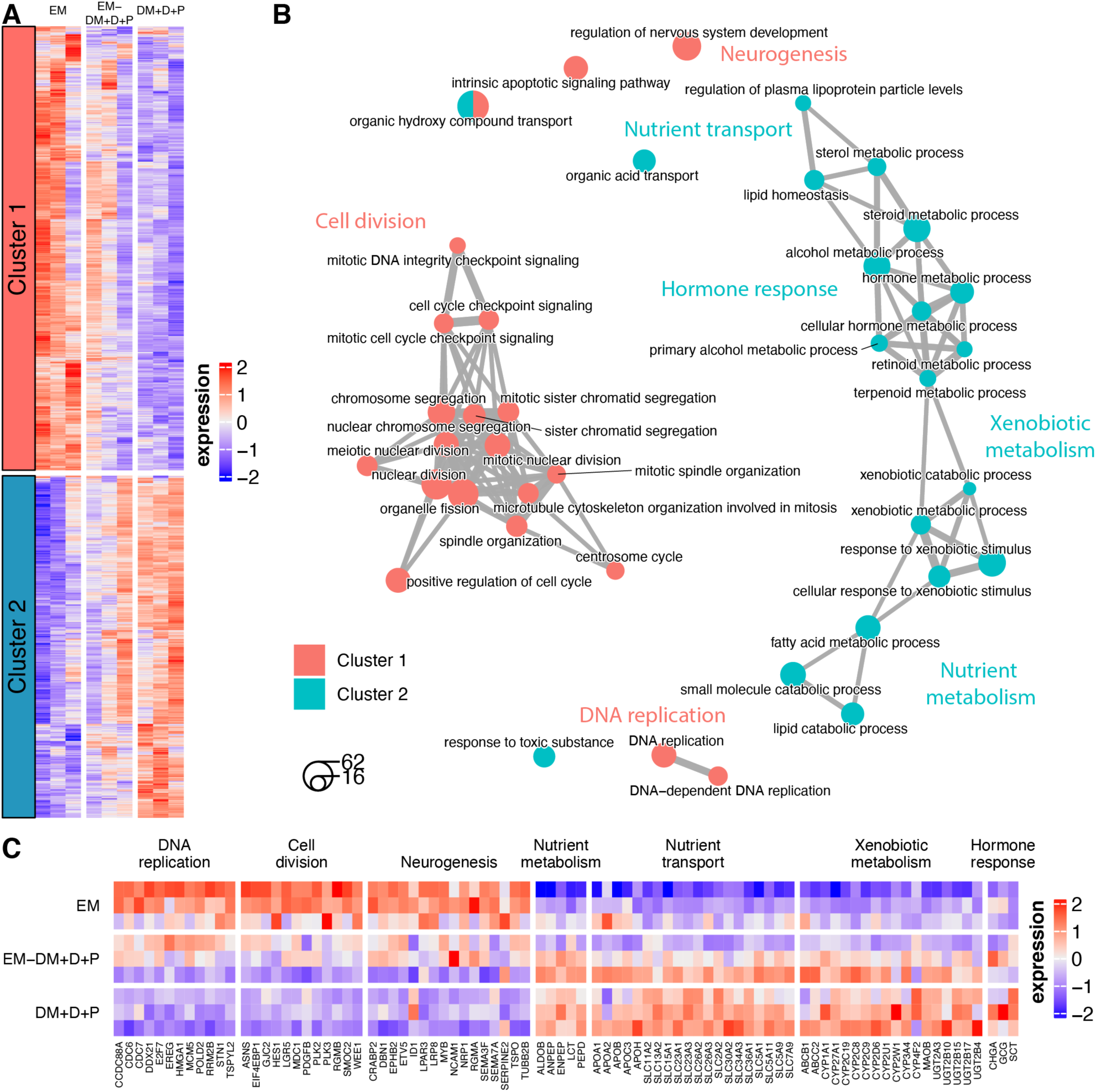
Balancing transit-amplifying/stem cell-specific and enterocyte- and enteroendocrine-specific functions in the EM-DM+D+P condition in Transwell systems. (A) Heatmap of average expression of the DEGs between media conditions in monolayers in Transwell systems, clustered in groups of genes with similar expression profiles. Color scale represents row z-score. Columns correspond to donors, left=1, middle=2, right=3. (B) Network of enriched processes identified using the Gene Ontology: biological process database (top 20 with lowest adjusted p-value per cluster). Nodes depict enriched biological processes. Node size indicates the number of DEGs associated to the process. Edge thickness correlates with the number of DEGs shared between processes. (C) Heatmap of average expression of DEGs selected from enriched processes described in B. Color scale represents column z-score. Rows correspond to donors, upper=1, middle=2, lower=3.

Altogether, these genes and processes are signatures of the enrichment of transit-amplifying cells and intestinal stem cells in the EM condition and the enrichment of mature enterocytes and enteroendocrine cells in the DM+D+P condition in both intestinal organoids and Transwell systems. In contrast to organoids, Transwell systems allow for gradient conditions that balance proliferative and mature epithelial cell types and processes.

### Enrichment of intestinal functions and enhanced maturation of hiPSC-derived intestinal epithelial cells grown in intestine-on-chip relative to organoids and Transwells

Lastly, we wanted to investigate the biological processes regulated by the culture environment of conventionally used intestinal model systems. We recently demonstrated that exposure to the EM-DM+D+P condition in a microfluidic hiPSC-derived intestine-on-chip system induces an epithelial composition that resembles the human small intestine^15^. Here, we compared the transcriptomic profiles of hiPSC-derived intestinal epithelial cells grown as organoids (exposed to DM+D+P), in Transwell systems and in a commercially available intestine-on-chip system (both exposed to EM-DM+D+P) (Figures 5A, S1C). Pairwise analysis between the model systems and subsequent clustering of DEGs resulted in three clusters that reflect different expression profiles between the systems (Figures 5B, Data S1). The DEGs upregulated in intestinal organoids (cluster 1) encompass multiple genes related to mineral and metal metabolism, primarily metallothionein genes (e.g. *MT1F, MT1X* and *MT1H*)^26^ (Figures 5B-D). The DEGs upregulated in Transwell systems (cluster 3) are related to cell division, suggesting that even though the Transwell and intestine-on-chip systems were both exposed to the EM-DM+D+P condition, the microenvironment of the intestine-on-chip might accelerate epithelial differentiation (Figures 5B-D). In addition, processes related to extracellular matrix organization (including genes *TGFB1, TGFB2, COL2A1, COL4A6, LAMB1, LAMB2* and *ACTA2*) and neurogenesis (e.g. *TUBB2B, NCAM1* and *RGMA*) are enriched in Transwell systems ^27–29^ (Figures 5B-D). Accordingly, we observed more VIM-expressing epithelial cells in Transwell systems compared to organoids and intestine-on-chip systems, suggesting a potential epithelial-to-mesenchymal transition, as observed in a previous study^15^ (Figure S1D). Genes specifically upregulated in intestine-on-chip systems (cluster 2) primarily included enterocyte-associated genes related to small intestinal functions such as nutrient metabolism (e.g. *ANPEP, SI, LCT, MGAM, PEPD, ALPI, LIPA* and *TMPRSS15*), nutrient transport (e.g. *SLC2A2, SLC2A5, SLC15A1, SLC26A3, APOA1, APOB* and *FABP2*) and xenobiotic metabolism (e.g. *CYP3A4, CYP3A5, CYP3A7, CYP1A1, CYP2C9, CYP2C18, CYP2D6, CYP2J2, UGT1A1, MAOB, ABCC2* and *ABCB1*)^16,17,19,20^ (Figures 5B-D). Strikingly, DEGs in clusters 1 (enhanced in organoids) and 3 (enhanced in Transwell systems) were enriched for processes related to embryonic development of the epithelium and differentiation of non-intestinal tissues (Figures 5B-D). The expression of homeobox genes (e.g. *HOXA9, HOXB2, HOXB4* and *HOXA1*) and genes involved in hedgehog signaling (e.g. *SHH, IHH, SMO, HES1* and *SUFU*), instrumental for patterning differentiation of tissues in mammalian embryos^30,31^, contributed to the identification of these processes (Figure 5D). This suggests that the epithelial tissue of organoids and in Transwell systems display a more fetal phenotype and resemble an earlier state of intestinal development compared to the tissue in intestine-on-chip systems.

**Figure 5.**
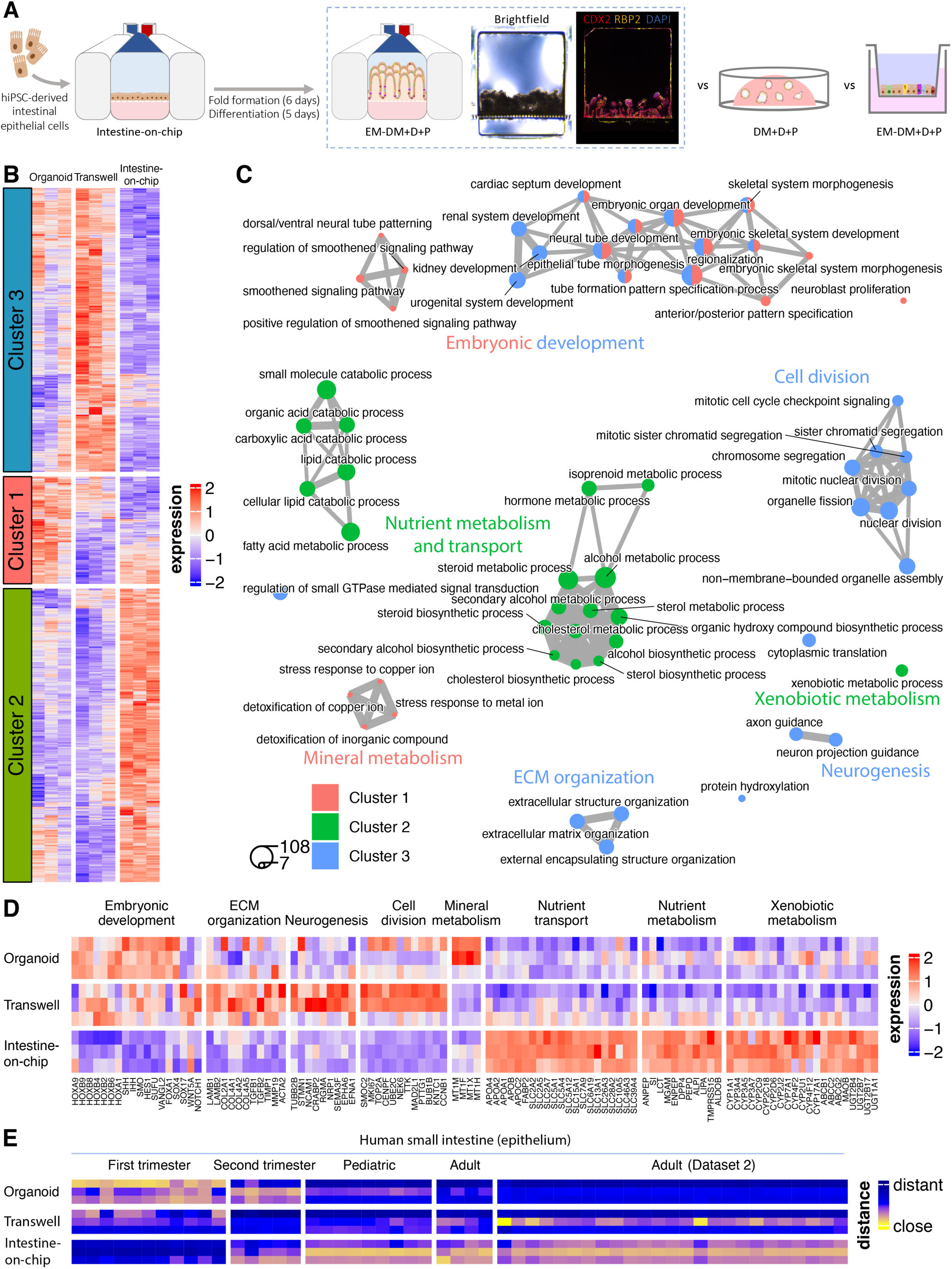
Enrichment of intestinal functions and maturation of epithelial cells grown in an intestine-on-chip relative to organoids and Transwell systems. (A) Schematic overview of the experimental set-up. Representative brightfield and immunofluorescent confocal image of cross-sectional slices of the intestine-on-chip system exposed to the EM-DM+D+P condition stained for markers characteristic of intestinal epithelial cells (CDX2) and enterocytes (RBP2). hiPSC = human induced pluripotent stem cell, EM = expansion medium, DM+D+P = differentiation medium + DAPT + PD0325901. (B) Heatmap of average expression of the DEGs between organoids (DM+D+P), Transwell and intestine-on-chip systems (both EM-DM+D+P), clustered in groups of genes with similar expression profiles. Color scale represents row z-score. Columns correspond to donors, left=1, middle=2, right=3. (C) Network of enriched processes identified using the Gene Ontology: biological process database (top 20 with lowest adjusted p-value per cluster). Nodes depict enriched biological processes. Node size indicates the number of DEGs associated to the process. Edge thickness correlates with the number of DEGs shared between processes. ECM = extracellular matrix. (D) Heatmap of average expression of DEGs selected from enriched processes described in C. Color scale represents column z-score. Rows correspond to donors, upper=1, middle=2, lower=3. (E) Scaled distance between epithelial cells of the human small intestine and hiPSC-derived intestinal epithelial cells as organoids, in Transwell systems and in intestine-on-chip systems. Data for the different age groups of the human small intestine are derived from the Gut Cell Atlas^28,32^ and Ramírez-Sánchez et al.^33^ (dataset 2). Distances are generated based on DEGs between the epithelial compartment and the mesenchymal, neural, endothelial and immune compartments in the data from the Gut Cell Atlas. Smaller distance between samples (yellow values) indicates higher similarity and greater distance (blue values) indicates lower similarity. Rows correspond to donors as in D.

To test whether the intestine-on-chip promotes intestinal epithelial maturation, we compared the transcriptome data of the organoids, Transwell and intestine-on-chip systems to publicly available transcriptome data from human small intestinal epithelial cells at different developmental stages from the Gut Cell Atlas^28,32^ and Ramírez-Sánchez et al.^33^. Indeed, organoids displayed the smallest distance and thus the highest resemblance to the fetal small intestine, as determined by a distance analysis based on genes characteristic of human small intestinal epithelial cells (Figure 5E). Transwell systems did not show a specific resemblance to any intestinal developmental stage, while the intestine-on-chip systems showed the highest and most specific similarity to pediatric and adult human small intestinal expression profiles (Figure 5E).

According to this data, the intestine-on-chip drives the gene expression profile of hiPSC-derived intestinal epithelial cells toward processes related to nutrient and xenobiotic metabolism and more mature stages of small intestinal development (pediatric and adult), whereas organoids and Transwell systems may provide a better representation of fetal intestinal processes, including embryonic epithelial patterning, and specific processes such as mineral metabolism (organoids) and non-epithelial processes such as neurogenesis (Transwell systems).

## Discussion

Many studies have investigated the effect of *in vitro* culture conditions on ASC-derived intestinal epithelial cells, but only limited data is available for hiPSC-derived cells^7–10,14,34^. In this study, we investigated the controlled induction of hiPSC-derived intestinal epithelial subtypes and the biological processes induced by the culture microenvironment of organoids, Transwell and intestine-on-chip systems. We found that hiPSC-derived intestinal epithelial cells can be steered toward specific lineages using similar growth factors as have been described for ASC-derived intestinal epithelial cells^10,14^. Activation of the WNT pathway and inhibition of the BMP pathway maintained proliferating epithelial cells in the EM condition. A 5-day exposure to the DM condition, which excluded these activators and inhibitors, resulted in a near absence of MKI67-positive dividing cells and an induction of RBP2-positive enterocytes and MUC2-positive goblet cells. The introduction of a NOTCH inhibitor (DAPT) to the DM condition (DM+D) further induced MUC2-positive goblet cells. Additional MAPK inhibition by exposure to PD0325901 (DM+D+P) was necessary to induce low levels of CHGA-positive enteroendocrine cells (mainly expressing hormones *MLN*, *GCG* and *SCT*) and reduced the MUC2-positive goblet cells. Overall, these principles can be applied to steer hiPSC-derived intestinal tissues toward a desired epithelial composition, regardless of the model system used. To balance epithelial proliferation and differentiation, and thus increase epithelial diversity and physiological relevance, basolateral exposure to the EM condition and apical exposure to a type of DM can be applied in both Transwell and intestine-on-chip systems^15^. The fact that this works in the static and dynamic environment of both systems demonstrates that continuous renewal of medium in a microfluidic system is not required. The levels of CHGA-positive enteroendocrine cells induced in the DM+D+P condition (below 1%) are lower than previously reported by using MAPK inhibition in human ASC-derived intestinal organoids (10-20%)^35,36^ but fall in the range reported for the human intestine (0-1%)^17,37^. The levels of MUC2-positive goblet cells induced in the DM and DM+D conditions (20-30%) resemble the abundance of goblet cells in the human large intestine (∼20%), while the DM+D+P condition results in levels around 10%, closer to that of the small intestine (∼5%)^17,37^.

When differentiating hiPSCs to intestinal cells, mesenchymal cells co-develop and expand in EM conditions^4,15^. In the current study we aimed to investigate hiPSC-derived intestinal epithelial cells specifically and set out to generate intestinal tissues with negligible numbers of mesenchymal cells through selection of EPCAM-positive cells and consecutive rounds of single-cell passaging. Nevertheless, we still identified some cells expressing the mesenchymal marker VIM and observed that VIM expression levels correlated positively with the expression of cell cycle, stem cell and Paneth cell genes and negatively with epithelial differentiation in the different donors in the Transwell system. Moreover, the intestine-on-chip exposed to EM-DM+D+P in this study had a lower expression of genes associated with the cell cycle and Paneth cells and a lower tissue height when compared to an intestine-on-chip in the same medium condition that included mesenchymal and neural cells that we described in a previous study^15^. This data suggests that hiPSC-derived epithelial cells may depend on mesenchymal-derived factors to sustain a proliferative and potentially stem cell- and Paneth-like state. Inclusion of mesenchymal cells or addition of mesenchymal-derived factors (such as WNT ligands or R-spondins) to the EM condition may help facilitate the long-term maintenance of dividing hiPSC-derived intestinal epithelial cells.

As shown here, the unique culture environments of model systems change the cellular and gene expression profiles of epithelial cells, likely due to differences such as the physiological shear stress induced by continuous fluid flow, stiffness profiles induced by embedding in extracellular matrix or the configuration into multiple compartments, which enables dual exposure to EM and DM. The cells in the Transwell system showed an increased expression of neuron-associated and extracellular matrix genes and had a higher expression of epithelial cells expressing VIM. This might indicate that the Transwell microenvironment or the longer culturing times required to establish a polarized epithelial barrier in this static environment, enhance the presence of mesenchymal and neural cells and phenotypes resembling epithelial-to-mesenchymal transition, as we observed earlier in intestine-on-chip systems^15^. Intestinal organoids displayed transcriptional profiles indicating an upregulation of mineral absorption and metallothioneins, which protect against metal toxicity in the human intestine^26^. Potentially, this results from an enhanced induction of metallothionein-expressing enterocytes, which co-develop along with more classical enterocytes in hiPSC-derived intestinal systems^15^.

Although we could induce expression profiles reflecting mature enterocyte functions in organoids and Transwell systems upon exposure to (EM-)DM+D+P, they were further enhanced in the dynamic environment of the intestine-on-chip. Diverse genes encoding digestive enzymes (e.g. *SI, LCT, ANPEP, LIPA* and *ALDOB*) and transporters of the SLC- and apolipoprotein (APO) family (e.g. *SLC2A2, SLC2A5, SLC15A1, SLC13A1, APOA4, APOA1* and *APOB*) involved in digestion and transport of carbohydrates, proteins, lipids and ions showed increased expression in the intestine-on-chip compared to organoids and Transwell systems^16,17^. Moreover, enzymes of the Cytochrome P450 family involved in the conversion of therapeutic compounds in the human intestine (e.g. *CYP3A4, CYP3A5, CYP3A7, CYP2C9, CYP2D6* and *CYP2J2*) were upregulated and included genes of the CYP3A and CYP2C families that represent 80% and 18% of the intestinal Cytochrome P450 genes, respectively^38^. These processes and genes cover well-established metabolic processes in the human small intestine and include genes that display high inter-individual genetic variability (e.g. lactase (*LCT*) and *CYP2D6*)^1,39^, emphasizing the value of hiPSC-derived intestinal epithelial cells for personalized modeling of intestinal digestion and drug metabolism upon exposure to relevant culture conditions. The elevated expression of nutrient and drug metabolism upon exposure to continuous fluid flow was in concordance with findings from studies using human ASC-derived intestinal epithelial cells: genes associated with lipid, protein and xenobiotic metabolism were upregulated in a human duodenum-derived intestine-on-chip when compared to organoids^9^ and in monolayers of duodenal epithelial cells after exposure to rotational flow profiles^13^. Moreover, an ASC-derived colon intestine-on-chip showed upregulation of CYP3A4 when compared to organoids^8^. In addition to enhanced enterocyte functions, microfluidic intestine-on-chip systems show potential to study nutrient and drug metabolism and absorption given their easy access to the basolateral and apical side of the epithelial barrier, enlarged epithelial surface area due to the presence of villus-like folds, physiological barrier integrity^15^ and ability to sustain dividing and differentiated epithelial subtypes. Moreover, they confer a unique opportunity to study the bioavailability of compounds after oral intake and first-pass metabolism when coupled to a liver-on-chip containing the same genetic background.

The relative fetal state of hiPSC-derived intestinal organoids was previously improved by a 6 to 12-week transplantation in mice^11,12,40^. Our study shows that maturation and specific intestinal processes can also be induced by exposure to DM-type conditions. Importantly, we demonstrate that the more physiological environment in the intestine-on-chip can improve the overall maturation state of hiPSC-derived intestinal epithelial cells *in vitro*. The gene expression profiles in the intestine-on-chip showed more similarity to later developmental stages of the human small intestine (pediatric and adult) than to fetal stages, whereas the profiles of organoids resembled fetal stages and the profiles in Transwell systems did not resemble any specific stage. Indeed, genes involved in embryonic developmental processes, particularly homeobox genes and genes involved in hedgehog signaling, were downregulated in intestine-on-chip systems when compared to organoids and Transwell systems. The enhanced maturation in the intestine-on-chip system was also achieved with a shorter culturing time of 11 days, as compared to 19 days in Transwell systems and 12 days as organoids.

In conclusion, exposure to activators and inhibitors of the WNT, BMP, NOTCH and MAPK pathways allows the controlled induction of hiPSC-derived intestinal epithelial lineages. While hiPSC-derived intestinal epithelial cells grown as organoids or in Transwell systems might represent fetal or developmental intestinal processes, intestine-on-chip systems could be more relevant for modeling pediatric or adult intestinal phenotypes and digestive processes of the human small intestine in a personalized manner.

## Acknowledgements

This work was supported by the Netherlands Organ-on-Chip Initiative, an NWO Gravitation project (024.003.001) funded by the Ministry of Education, Culture, and Science of the government of the Netherlands (R.Moerkens, C.W., S.W.). J.M is supported by a PhD scholarship from the Graduate School of Medical Sciences, University of Groningen. I.H.J. is supported by a Rosalind Franklin Fellowship from the University of Groningen and an NWO VIDI grant (016.171.047). We thank Kate Mc Intyre for editing the manuscript. We thank the iPSC/CRISPR facility, UMCG Microscopy and Imaging Center and Flow Cytometry Unit of the University Medical Center Groningen for their support and services. We thank Emulate Inc (Boston, USA) for kindly providing the Human Emulation System. We thank the members of the Netherlands Organ-on-Chip Initiative for insightful discussions and support regarding the implementation of organ-on-chip technology.

## Author contributions

R.Moerkens and J.M. designed and executed experiments. R.Moerkens analyzed experimental data and wrote the manuscript. M.B. analyzed RNA-sequencing data. E.S. and R.Modderman assisted in executing experiments. R.Moerkens, J.M., A.D.R., J.P., C.P. and R.J.B. (co-)developed and/or were consulted for methodologies. C.W. assisted in conceiving the project. I.H.J. and S.W. supervised the project and provided feedback on experiment design, data analysis and writing.

## Methods

### Resource availability

#### Lead contact

Further information and requests should be directed to the lead contact, Sebo Withoff.

#### Materials availability

This study did not generate new unique reagents.

#### Data and code availability

- The RNA-sequencing data will be shared by the lead contact upon request.
- The code required to reproduce the RNA-sequencing analysis detailed above can be found at https://github.com/umcg-immunogenetics/Organoid_Transwell_Chip_Moerkens_2024.
- Any additional information required to reanalyze the data reported in this paper is available from the lead contact upon request.

### Experimental model and study participant details

#### Cell lines and culturing conditions

Human induced pluripotent stem cell (hiPSC) lines were generated from urine-derived renal epithelial cells of three healthy donors (two male, one female) by the iPSC/CRISPR facility (ERIBA, UMCG, Groningen) using a lentiviral vector described earlier^41^. The expression of pluripotency markers and absence of differentiation markers was verified on protein and RNA level in the hiPSC lines. The hiPSC lines were maintained in mTeSR Plus (Stemcell Technologies, #05825) on plates coated with hESC-qualified Matrigel (Corning #354277). Passaging was performed every 3-4 days using ReLeSR (Stemcell Technologies, #05872) according to manufacturer’s instructions. Cells were cultured in a humidified environment, at 37 °C in 5% CO2. hiPSC lines were cryopreserved as fragments in CryoStor CS10 (Stemcell Technologies #7930). The experiments with hiPSC lines were approved by the ethics committee of the University Medical Centre Groningen, document no. METC 2013/440 and written consent was obtained from the donors.

### Method details

#### Differentiation of hiPSCs to intestinal organoids

The differentiation procedure to generate intestinal organoids from hiPSCs was adapted from previously described protocols^34,42^. Briefly, hiPSCs were grown to ∼75-80% confluence in a 24-well plate. Definitive endoderm was induced by 3-day exposure to RPMI 1640 (ThermoFisher #11875093) supplemented with L-glutamine (2 mM, ThermoFisher #25030081), Penicillin-Streptomycin (100 Units/ml; 100 μg/ml respectively, ThermoFisher # 15140122), Activin A (100 ng/ml, R&D Systems, #338-AC) and an increasing concentration of defined FBS (day 1: 0%, day 2: 0.2%, day 3: 2% vol/vol, GElifesciences, Hyclone #SH30070.02). Additionally, Wnt3a was supplemented only on day 1 (25 ng/ml, R&D Systems, #5036-WN/CF). Thereafter, mid/hindgut was induced by 4-day exposure to Advanced DMEM/F12 (ThermoFisher #12634010) supplemented with L-glutamine (2 mM, ThermoFisher #25030081), Penicillin-Streptomycin (100 Units/ml; 100 μg/ml respectively, ThermoFisher #15140122), FGF4 (500 ng/ml, R&D Systems, #235-F4/CF), CHIR99021 (3 μM, Tocris, #4423) and defined FBS (2% vol/vol, GElifesciences, Hyclone #SH30070.02), which was refreshed daily. For the induction of human intestinal organoids, the adherent monolayer containing three-dimensional structures and the free-floating spheres that developed during mid/hindgut induction were mechanically harvested and fragmented by repeated pipetting. After gravity-based sedimentation of fragments, supernatant was discarded and the fragments were re-suspended in Basement Membrane Matrigel (Corning #354234) and plated in domes in a 24-well plate (ThermoFisher #142475). After 10 minutes of incubation at 37°C, Matrigel domes had solidified and were overlaid with Advanced DMEM/F12 (ThermoFisher #12634010) supplemented with L-glutamine (2 mM, ThermoFisher #25030081), Penicillin-Streptomycin (100 Units/ml; 100 μg/ml respectively, ThermoFisher # 15140122), Noggin (100 ng/ml, R&D Systems, #6057-NG/CF), EGF (100 ng/ml, R&D Systems, #236-EG), CHIR99021 (2 μM, Tocris, #4423), B27 (1x, ThermoFisher #17504044). Medium was replaced every other day. After 7-10 days of culture, the density of the organoids was reduced by passaging: Matrigel domes were mechanically dislodged and organoids were released from the domes by repeated pipetting. The suspension was spun down at 400xg for 3 minutes and the supernatant and Matrigel layer were aspirated. The organoids were washed twice in DPBS (Gibco, #14190-094), re-suspended in fresh Basement Membrane Matrigel and plated in domes as described before.

#### Selection of epithelial cells from intestinal organoids

The majority of mesenchymal cells that co-develop during the differentiation were removed to enrich intestinal epithelial cells after 17-18 days of growing the human intestinal organoids in Matrigel domes. Matrigel domes were mechanically dislodged and organoids were released from the domes by repeated pipetting. The suspension was spun down at 400xg for 3 minutes and the supernatant and Matrigel layer were aspirated. The organoids were washed twice in DPBS, re-suspended in warm TrypLE Select (ThermoFisher #12563029) and incubated for 5-6 minutes at 37 °C. The organoids were then dissociated by repeated pipetting. Depending on whether the organoids had sufficiently dissociated, the incubation with TrypLE Select was prolonged. DPBS supplemented with FCS (10% vol/vol, Gibco, #10270-106) was added to the cell suspension to stop TrypLE Select activity after the organoids were dissociated to a single-cell suspension, which was then passed through a 40-μm filter. The filtered cell suspension was spun down at 400xg for 3 minutes and re-suspended in DPBS supplemented with FCS (10% vol/vol). Epithelial cells were enriched using the EasySep™ Human EpCAM Positive Selection Kit II (Stemcell Technologies, #17846) according to manufacturer’s instructions. The resulting epithelial cells were counted and cryopreserved in CryoStor CS10 (Stemcell Technologies, #7930).

#### Expansion of intestinal epithelial cells

Intestinal epithelial cells were thawed, resuspended in Basement Membrane Matrigel and plated in domes as described before at a concentration of 1.000 cells/μl Matrigel. Solidified domes were overlaid with Advanced DMEM/F12 (ThermoFisher #12634010) supplemented with L-glutamine (2 mM, ThermoFisher #25030081), Penicillin-Streptomycin (100 Units/ml; 100 μg/ml respectively, ThermoFisher # 15140122), Noggin (100 ng/ml, R&D Systems, #6057-NG/CF), EGF (100 ng/ml, R&D Systems, #236-EG), CHIR99021 (2 μM, Tocris, #4423), B27 (1x, ThermoFisher #17504044), SB202190 (10 µM, Tocris #1264/10), A83-01 (500 nM, Tocris #2939/10), hereafter named ‘expansion medium’ (EM), and Y-27632 (10 µM, Tocris #1254/10). After 48 hours, the medium was replaced with fresh EM to remove Y-27632 and EM was replaced every other day. For 4-5 weeks, human intestinal organoids were grown and passaged weekly as fragments or single cells. For fragment passaging, culture medium was replaced with cold Advanced DMEM/F12, Matrigel domes were mechanically dislodged and organoids were released from the domes by repeated pipetting. The suspension was spun down at 400xg for 5 minutes at 4°C, the Matrigel layer was discarded and organoids were resuspended in cold Advanced DMEM/F12 and fragmented by repeated pipetting using a P10 tip on top of a P1000 tip. The suspension was spun down at 400xg for 5 minutes at 4°C and the fragments were resuspended in Basement Membrane Matrigel and plated in domes overlaid with EM as described before. For single cell passaging, the same procedure was followed as for fragment passaging, but fragments were re-suspended in warm TrypLE Select (ThermoFisher #12563029), dissociated to a single cell suspension as described before and passed through a 70-μm filter. The filtered cell suspension was spun down at 400xg for 5 minutes, resuspended in Basement Membrane Matrigel at a concentration of 1.000 cells/μl Matrigel and plated in domes overlaid with EM containing Y-27632 (10 µM, Tocris #1254/10) as described before. After 48 hours, EM was replaced to remove Y-27632.

#### Seeding and maintenance of intestinal epithelial cells as organoids, in Transwells and in intestine-on-chips

To seed organoids, intestinal epithelial cells were resuspended as single cells in Basement Membrane Matrigel at a concentration of 500 cells/μl Matrigel and plated in domes overlaid with EM containing Y-27632 (10 µM, Tocris #1254/10) as described before. Domes were plated in 24-well plates for RNA harvesting and flow cytometry or in Nunc™ Lab-Tek™ Chamber Slide Systems (ThermoFisher #178599PK) and spread over the surface of the well before solidification for immunofluorescent microscopy. To seed Transwell systems (Corning #3401), inserts were coated with Basement Membrane Matrigel (83 µg/ml, Corning #354234) diluted in Advanced DMEM/F12 (ThermoFisher #12634010) and incubated for 1 hour at room temperature. EM containing Y-27632 (10 µM, Tocris #1254/10) was introduced to the bottom reservoir and the Matrigel suspension was aspirated from inserts. Intestinal epithelial cells were resuspended as single cells in EM containing Y-27632 (10 µM, Tocris #1254/10) and 2×10^5^ cells were seeded per insert. Intestine-on-chips (PDMS-based Chip-S1, Emulate Inc Boston, MA) were seeded as described previously^15^. In short, the two microfluidic channels were activated, coated with Basement Membrane Matrigel (83 µg/ml, Corning #354234) diluted in Advanced DMEM/F12 (ThermoFisher #12634010) and 3×10^5^ intestinal epithelial cells suspended in EM containing Y-27632 (10 µM, Tocris #1254/10) were seeded in the top channel of each chip. The chips were incubated for 3 hours in a humidified environment at 37 °C in 5% CO2 until cells were attached to the membrane, the top channel of the chip was gently washed and chips were connected to the Emulate instrument. The flow rate of the media within both channels was 30 ul/hour, which imposes a wall shear stress on the cells of 0.05 mPa in the top channel and 1.0 mPa in the bottom channel. Culture medium added to the chips was always first equilibrated. Cells were grown in EM for 6 days (intestine-on-chip), 7 days (organoids) or 14 days (Transwell) and thereafter exposed to different medium conditions for 5 days: EM, DM, DM+D or DM+D+P. DM was composed of Advanced DMEM/F12 (ThermoFisher #12634010) supplemented with L-glutamine (2 mM, ThermoFisher #25030081), Penicillin-Streptomycin (100 Units/ml; 100 μg/ml respectively, ThermoFisher #15140122), EGF (100 ng/ml, R&D Systems, #236-EG), B27 (1x, ThermoFisher #17504044). DM+D was composed of DM including DAPT (10 µM, Tocris #2634). DM+D+P was composed of DM+D including PD0325901 (0.1 µM, Tocris #4192). The concentration of dissolvent was equal in all medium conditions. All systems were maintained in a humidified environment at 37 °C in 5% CO2 and respective media were refreshed every 2 days for organoids and Transwell systems and using continuous medium flow in the intestine-on-chip systems.

#### Immunofluorescent microscopy

Intestine-on-chip systems were cut in half perpendicular to the channels using a microtome blade (pfm medical #207500002) and half of each chip was used for immunofluorescent microscopy. Intestinal epithelial cells grown as organoids, in Transwell systems and in intestine-on-chip systems were fixed to be used for immunofluorescent microscopy. The cells were washed twice with DPBS and incubated with paraformaldehyde (PFA, 4% vol/vol, ThermoFisher #28908) diluted in DPBS for 10 minutes (Transwell and intestine-on-chip systems) or 60 minutes (organoids) at room temperature. The cells were washed twice with DPBS and stored in DPBS at 4 °C until immunofluorescent staining. For intestine-on-chip systems, thin sections of the PDMS were cut away on all sides of the chip using a microtome blade (pfm medical #207500002). Cross-sectional slices (200 µm thick) were generated while chips were submerged in DPBS with ice using a vibratome (VT1000S, Leica; speed: 0.075 mm/s, frequency: 60-65 Hz, blade angle: 5°, razorblades: Astra Superior Platinum Double Edge). The lower part of the blade holder of the vibratome was replaced by a manually designed piece with a flatter design, to prevent collision of this piece with the chip while sectioning. For Transwell systems, inserts were removed from the holder using a scalpel and divided in quarters using a microtome blade (pfm medical #207500002). The organoids embedded in Matrigel domes, tissue on Transwell inserts and in intestine-on-chip slices were permeabilized by incubation in Triton X-100 (0.1% vol/vol, Sigma Aldrich #T8787) diluted in DPBS for 10-15 minutes (Transwell systems) or 30 minutes (organoids and intestine-on-chip systems) at room temperature and blocked by incubation in bovine serum albumin (BSA, 3% weight/vol, Sigma Aldrich #A2153) diluted in DPBS for 1 hour at room temperature. Samples were stained overnight at 4 °C in BSA (3% weight/vol) diluted in DPBS containing the primary antibodies. The next day, samples were washed twice with DPBS and stained in BSA (3% weight/vol) diluted in DPBS containing the secondary antibodies for 2 hours at room temperature in the dark. Then samples were washed three times with DPBS and Transwell inserts and intestine-on-chip slices were mounted on a glass microscope slide in mounting medium with DAPI (Vector Laboratories #H-2000), while Matrigel domes containing organoids were submerged in mounting medium with DAPI. Samples were kept at 4 °C in the dark until imaging. Images were taken using a Leica SP8 CLSM confocal immunofluorescent microscope (using the 10x or 20x objective). Results were analyzed using the Leica LAS X software. Primary and secondary antibodies are listed in Tables S1 and S2.

#### Dissociation of single cells from intestinal organoids, Transwell and intestine-on-chip systems

Intestine-on-chip systems were cut in half perpendicular to the channels using a microtome blade (pfm medical #207500002) and half of each chip was used for dissociation. Intestinal epithelial cells grown as organoids, in Transwell systems and in intestine-on-chip systems were dissociated to single cells to be used for flow cytometry analysis. Organoids were released from Matrigel domes and dissociated using TrypLE Select as described before. Transwell and intestine-on-chip systems were washed twice with DPBS, warm TrypLE Select was introduced to the bottom and top compartments and systems were incubated at 37 °C. Every 10 minutes, cells were detached by repeated pipetting and fresh TrypLE Select was introduced. The cell suspension was collected in a microcentrifuge tube, incubated for 5 minutes at 37 °C and further dissociated to a single cell suspension by gentle repeated pipetting. The single-cell suspension was added to a centrifuge tube containing Advanced DMEM/F12 (ThermoFisher #12634010) supplemented with FCS (10% vol/vol) and Y-27632 (10 µM) and stored on ice. Previous steps were repeated 3-4 times until the tissue was removed from the insert of the Transwell system and the top channel of the intestine-on-chip. The resulting cell suspension was passed through a 70-μm filter, centrifuged (400xg, 5 minutes, 4 °C) and resuspended in Advanced DMEM/F12 (ThermoFisher #12634010) supplemented with FCS (10% vol/vol) and Y-27632 (10 µM).

#### Flow cytometry

For each model system, 1×10^5^ dissociated cells per condition were used in flow cytometry analysis. A fraction of the samples of each system was pooled to be used as ‘unstained sample’ and ‘single-antibody stained samples’. The remainder of the cells were stained using the Zombie Aqua Fixable Viability Kit (BioLegend #423101) according to the manufacturer’s instructions. Subsequently, the unstained and Zombie Aqua-stained samples were fixed by incubation in PFA (4% vol/vol, ThermoFisher #28908) diluted in DPBS for 10 minutes at room temperature, washed with DPBS and stored in DPBS with FCS (2% vol/vol) at 4 °C until flow cytometry analysis. For immunofluorescent staining, the cells were permeabilized and labelled with fluorophore-conjugated antibodies using the BD Perm/Wash buffer (BD Biosciences #554723) according to the manufacturer’s instructions. BD Horizon Brilliant Stain Buffer (BD Biosciences #563794) was included in the antibody mixture in an equal volume to the total volume of the antibodies. The cells were resuspended in DPBS with FCS (2% vol/vol) and analyzed using the Cytek Aurora. Results were analyzed using the Kaluza Analysis software (Beckman Coulter Life Sciences). Model system-specific unstained and ‘fluorescence-minus-one’ samples were included in the analysis as the basis for the gating strategy. Single-antibody stained samples were used for compensation. Antibodies are listed in Table S3.

#### RNA isolation and sequencing

RNA was isolated from intestinal epithelial cells grown as organoids, in Transwell systems and in intestine-on-chip systems to be used for RNA sequencing. For organoids, culture medium was replaced with cold Advanced DMEM/F12, Matrigel domes were mechanically dislodged and organoids were released from the domes by repeated pipetting. The suspension was spun down at 400xg for 5 minutes at 4°C, the Matrigel layer was discarded and organoids were washed in DPBS. Organoids were resuspended in Lysis/Binding Buffer from the mirVana™ miRNA Isolation Kit (ThermoFisher #AM1561) and lysed by repeated pipetting. For Transwell systems, cells were washed twice with cold DPBS, Lysis/Binding Buffer was added to the insert and cells were lysed by repeated pipetting. For the intestine-on-chip systems, cells were dissociated from the top channel as described before, washed twice with DPBS and lysed by repeated pipetting in Lysis/Binding Buffer. The cell lysates of all model systems were transferred to an RNase-free microcentrifuge tube, vortexed for a few seconds to obtain a homogeneous lysate and stored at -80°C until RNA isolation. RNA was isolated using the mirVana™ miRNA Isolation Kit (ThermoFisher #AM1561) according to manufacturer’s instructions. High RNA integrity (RIN > 8.3) was confirmed using the Agilent RNA ScreenTape (#5067-5576) on the Agilent TapeStation 4200. RNA library preparation and sequencing were performed in one batch at BGI Tech Solutions (Hong Kong) according to the ‘DNBSEQ Eukaryotic Strand-specific mRNA library’ protocol. Briefly, oligo-dT beads were used to isolate mRNA from total RNA. mRNA molecules were fragmented and cDNA was synthesized. Sequencing adapters were ligated to cDNA fragments and PCR amplification was performed. Library quality and yield were determined and sequencing was performed using a DNBseq sequencing platform with a 150-bp paired-end (PE150) kit.

#### Gene expression quantification and quality control

Adapter sequences and low-quality reads were removed using the SOAPnuke software^43^ developed by BGI Tech Solutions (Hong Kong). The trimmed fastQ files were aligned to humanG1Kv37 reference genome (Ensembl Release 75) using Hisat (version 0.1.5)^44^ with default settings, and aligned reads were sorted using SAMtools (version 1.2)^45^. Gene-level quantification was performed using HTSeq-count HTSeq (version 0.6.1p1)^46^ with -- mode=union and Ensembl version 75 as gene annotation database. Quality control metrics were calculated for the raw sequencing data using FastQC (version 0.11.3)^47^ and for the aligned reads using Picard-tools (version 1.130)^48^. GATK tool SplitNCigarReads was used to split reads into exon segments and hard-clip sequences overhanging into the intronic regions. Variant calling was done using HaplotypeCaller in GVCF mode. All samples were then jointly genotyped by taking the gVCFs produced earlier and running GenotypeGVCFs to create a set of raw SNP and indel calls per chomosome^49^.

#### Differential gene expression analysis

The differentially expressed genes (DEGs) between different conditions were identified using the R package DEseq2 (version 1.44.0) including donor as covariate in the DE model. Genes were filtered to only those with 10 or more reads in at least 3 samples. Analysis was performed by multiple comparisons in the relevant subsets of the data, as follows: EM vs DM+D+P (organoids), EM vs DM+D+P and EM vs EM-DM+D+P and EM-DM+D+P vs DM+D+P (Transwell systems), organoid vs Transwell and organoid vs intestine-on-chip and Transwell vs intestine-on-chip (DM+D+P condition for organoids and EM-DM+D+P condition for Tranwell and intestine-on-chip systems). DEGs were filtered on having an absolute log2FoldChange ≥ 1 and an adjusted p-value < 0.01. Expression levels of DEGs were presented in Heatmaps as normalized or scaled counts and used for clustering and pathway enrichment analysis.

#### Clustering of DEGs and pathway enrichment analysis

The gene expression matrix was VST-normalised using DESeq2 (version 1.44.0) and filtered to include DEGs only. Normalized and scaled expression levels of DEGs were k-means clustered using the hclust() and dist() functions from R. Two clusters were generated for the organoid and Transwell medium condition comparisons and three clusters were generated for the model system comparison. The clustered DEGs were used to identify biological processes that were enriched in each cluster. This analysis was performed using the R package clusterProfiler (version 4.12.0) using the Gene Ontology: Biological Processes database. P-values were adjusted using the Benjamini-Hochberg procedure. The visualization of biological processes was done using the emapplot function from clusterProfiler, limiting the number of displayed processes to the top 20 per cluster based on adjusted p-value.

#### Distance matrix analysis

Genes representing the epithelial compartment of the human small intestine (annotated as ‘intestinal epithelial marker genes’) were determined using the single-cell RNA sequencing data of the Gut Cell Atlas^28,32^. The Gut Cell Atlas data was subsetted as follows: The inflammatory bowel disease-related samples (Diagnosis = ‘Pediatric Crohn Disease’) were excluded and cells of the small intestine were included (Region = ’SmallInt’). The subsetted data was used to identify DEGs using the FindMarkers function of Seurat (version 5.0.2), contrasting the intestinal epithelial cells (category = ’Epithelial’) to all other cellular compartments (category =’Mesenchymal’, ’Neuronal’, ’B cells’, ’Endothelial’, ’Myeloid’, ’Plasma cells’, ’Red blood cells’, ’T cells’). Intestinal epithelial marker genes were filtered as having adjusted p-value < 0.05 and falling in the fourth quantile of log2FoldChange (corresponding to log2FoldChange ≥ 2.7), resulting in 993 genes.

Pseudo-bulk data was generated of the single-cell RNA sequencing data of the Gut Cell Atlas^28,32^, subsetted for small intestinal epithelial cells using the same parameters as indicated above and additionally including epithelial cells only (category = ’Epithelial’) and excluding one donor (Sample.name = ’A32 (411C)’) based on the aberrant Paneth cell ratio of 50% from all epithelial cells in an ileal sample. Moreover, for individuals ‘T036’ and ‘T110’ several annotations were present in the data and the sample annotations describing the *EPCAM*-positive cells were used. To generate pseudo-bulk data, counts of the subsetted data were aggregated per age group (variable ‘Age_group’) and donor (variable ‘Sample.name’) using the Seurat function AggregateExpresssion. Raw counts of shared genes of the hiPSC-derived intestinal data (organoids, Transwell and intestine-on-chip system) described in this manuscript, the Gut Cell Atlas data and the RNA-sequencing data of the epithelial layer of human duodenal biopsies from healthy adults described by Ramírez-Sánchez et al.^33^ were merged in one count matrix. The euclidean distance between the samples was determined using the VST-normalized expression (DESeq2) of intestinal epithelial marker genes present in the count matrix using the dist() function in R and scaled per column for the visualization.

### Quantification and statistical analysis

The data was presented as median with interquartile range. Significant differences between the three media conditions were determined using a one-way analysis of variance (ANOVA) test with the Tukey multiple comparisons test. Differences between groups were considered statistically significant when P-value <0.05. Rstudio with the R package rstatix was used for statistical analysis^50^.

## Supplemental information

Document S1. Figure S1 and Tables S1-3

Data S1. Excel file containing the DEGs and the enriched biological processes generated in the RNA-sequencing analysis.

