## Supplementary material for "Intestine-on-chip enhances nutrient and drug metabolism and maturation of iPSC-derived intestinal epithelial cells relative to organoids and Transwells": Figure S1 and Tables S1-3

### Supplemental information

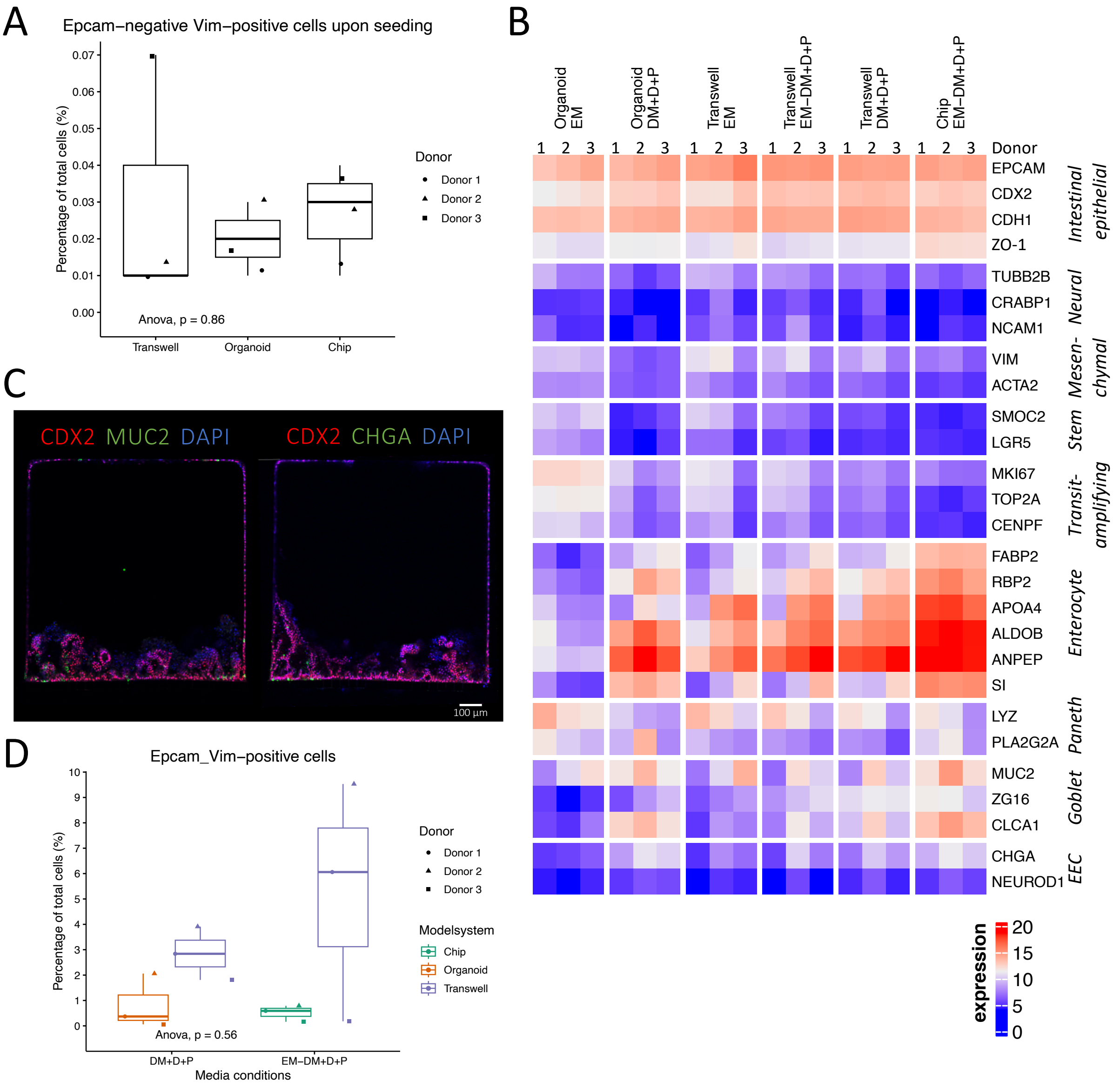


**Figure S1. Expression of epithelial-, mesenchymal- and neural-specific markers.** (A) Quantification of Epcam-negative VIM-positive cells present upon seeding of the model systems based on flow cytometry analysis, displayed as median values of three biological replicates. (B) Expression levels of genes associated with intestinal epithelial, mesenchymal and neural cells and multiple intestinal epithelial subtypes in diverse model systems and medium conditions. EEC = enteroendocrine cell. Color scale represents normalized counts. (C) Representative immunofluorescent confocal images of cross-sectional slices of hiPSC-derived intestinal epithelial cells in intestine-on-chip systems exposed to the EM-DM+D+P condition stained for markers characteristic of intestinal epithelial cells (CDX2), goblet cells (MUC2) and enteroendocrine cells (CHGA). (D) Quantification of Epcam-positive VIM-positive cells in different model systems based on flow cytometry analysis, displayed as median values of three biological replicates.

**Table S1. Primary antibodies used for immunofluorescent microscopy**

| Antigen | Dilution | Host/Isotype | Catalogue number | Manufacturer |
|---|---|---|---|---|
| CHGA | 1:100 | Mouse IgG1 | MA5-13096 | ThermoFisher |
| MUC2 | 1:200 | Mouse IgG1 | ab118964 | Abcam |
| LYZ | 1:200 | Mouse IgG2a | NB100-63062 | Novus Biologicals |

| | | | | |
|---|---|---|---|---|
| MKI67 | 1:200 | Rabbit IgG | ab16667 | Abcam |
| RBP2 | 1:100 | Rabbit polyclonal | HPA035866 | Sigma Aldrich |
| ZO-1 | 1:100 | Rabbit IgG polyclonal | 61-7300 | ThermoFisher |
| CDX2 | 1:100 | Goat IgG polyclonal | AF3665-SP | R&D systems |

**Table S2. Secondary antibodies used for immunofluorescent microscopy**

| Fluorophore | Dilution | Species reactivity | Catalogue number | Manufacturer |
|---|---|---|---|---|
| Alexa Fluor 488 | 1:250 | Mouse IgG (H+L) | 715-545-150 | Jackson ImmunoResearch |
| Alexa Fluor 647 | 1:250 | Goat IgG (H+L) | A-21447 | ThermoFisher |
| Cy3 | 1:250 | Rabbit IgG (H+L) | 711-165-152 | Jackson ImmunoResearch |

**Table S3. Antibodies used for flow cytometry**

| Antigen | Fluorophore | Dilution | Catalogue number | Manufacturer |
|---|---|---|---|---|
| VIM | Alexa Fluor 594 | 1:50 | 7675S | Cell Signaling Technology |
| EPCAM/CD326 | BUV737 | 1:100 | 748397 | BD Biosciences |
| CHGA | PE | 1:1000 | ab213341 | Abcam |
| MUC2 | PerCP | 1:67 | NBP2-34757PCP | Novus Biologicals |
| LYZ | Alexa Fluor 488 | 1:200 | NB100-63062AF488 | Novus Biologicals |
